# The Overlap Area as a Novel Measure of Effect Size in Neuroscience Research

**DOI:** 10.64898/2025.12.13.694093

**Authors:** Ying Wu, Hanan Woods, Kunhao Yuan, Digin Dominic, Zhen Qiu, Seth G.N. Grant

## Abstract

In experimental biomedical research, a common concern is whether a manipulation produces a biologically meaningful effect. Another concern is whether effect sizes are reliable, statistically significant, and generalizable. Traditional effect size measures, such as Cohen’s *d*, quantify mean differences but ignore variance heterogeneity between groups. This can result in biased effect size estimates and a lack of thresholds for statistical significance testing. Motivated by this, we introduce a novel effect size measure, termed overlap area (OA), which quantifies the difference between the population distributions of the experimental and control groups. A robust Bayesian method estimated OA, and random resampling determined OA thresholds for statistical significance in single and replicated experiments. Simulations confirmed the approach’s sensitivity and robustness. Applied to a real-world dataset, OA revealed that environmental enrichment affects the mouse brain synaptome. Moreover, we developed an open-source toolbox supporting OA as a powerful new measurement for conducting robust, reliable, and reproducible analyses of manipulation effects in neuroscience and related fields.

---

Effect size is a crucial statistical measure that enables researchers to quantify the magnitude of the effect of experimental manipulations. It is widely used and cited in psychology, neuroscience, and statistics. Since J. Cohen’s publications in 1988^1^ and 1992^2^, which have been cited over 20,000 times in psychology and statistics, effect size has been established as a fundamental component of experimental analysis. In neuroscience, Hentschke and Stüttgen (2011)^3^ introduced methods for computing effect sizes for neural data and, in psychology, Lakens (2013)^4^ provided a practical primer for calculating and reporting effect sizes in *t*-tests and analysis of variance. Effect size estimation has gained traction in neuroimaging and psychiatry, with studies by Reddan et al. (2017)^5^ and Szucs and Ioannidis (2017)^6^, who empirically assessed published effect sizes and statistical power in recent cognitive neuroscience and psychology literature. The study of effect size continues to evolve, with advances in software implementation^7^, methodological refinement^8^, and applied research^9,10^.

In most experimental research, scientists need not only to choose suitable effect size measures to assess whether a manipulation results in a biologically significant effect, but also to test the statistical significance of these effects in a single experiment. Additionally, establishing a clear significance testing threshold for replicated experiments is crucial to improving the reliability and reproducibility of research findings^11^. Two primary approaches have been proposed for assessing effect size in a single experiment. (1) Estimation, which focuses on calculating effect size measures such as Cohen’s *d* ^1,12,13^. This approach allows for the quantification of mean differences with groups. (2) Hypothesis testing, which evaluates the significance of an effect using methods such as the *t* -test ^14^ or the Bayes factor ^15^. However, Cohen’s *d* and *t*-test can be affected by small or unbalanced sample sizes, particularly when equal variances are assumed for the experimental and control groups. This can lead to biased estimates of effect size and obscure the true effect. Bayes factor hypothesis testing can be used in neuroscience only to quantify evidence of absence. These approaches measure mean differences, but they often overlook variance heterogeneity, lack intuitive interpretability of effect sizes, and provide limited guidance for establishing significance thresholds in replication studies.

To overcome these limitations, we introduce a novel measure termed overlap area (OA), along with a framework for its Bayesian estimation and hypothesis testing. First, we explain the issues related to existing methods, using Cohen’s *d* as an example. Next, using a series of simulation studies, we demonstrate how our proposed approach addresses effect size estimation and significance testing for both single and replicated experiments. The simulation results show that our method is robust and interpretable, overcoming limitations of traditional effect size measures. Finally, we applied this method to a real-world synaptic markers dataset, which successfully demonstrated that environmental manipulations induce effects across different brain regions. The developed toolbox, along with its functional description, further enhances the applicability of OA in neuroscience and related scientific disciplines.

## Results

### Workflow of the OA method

In the Bayesian framework, we introduced the OA, along with procedures for its estimation and hypothesis testing. This approach consists of five steps (Figure 1).

**Figure 1.**
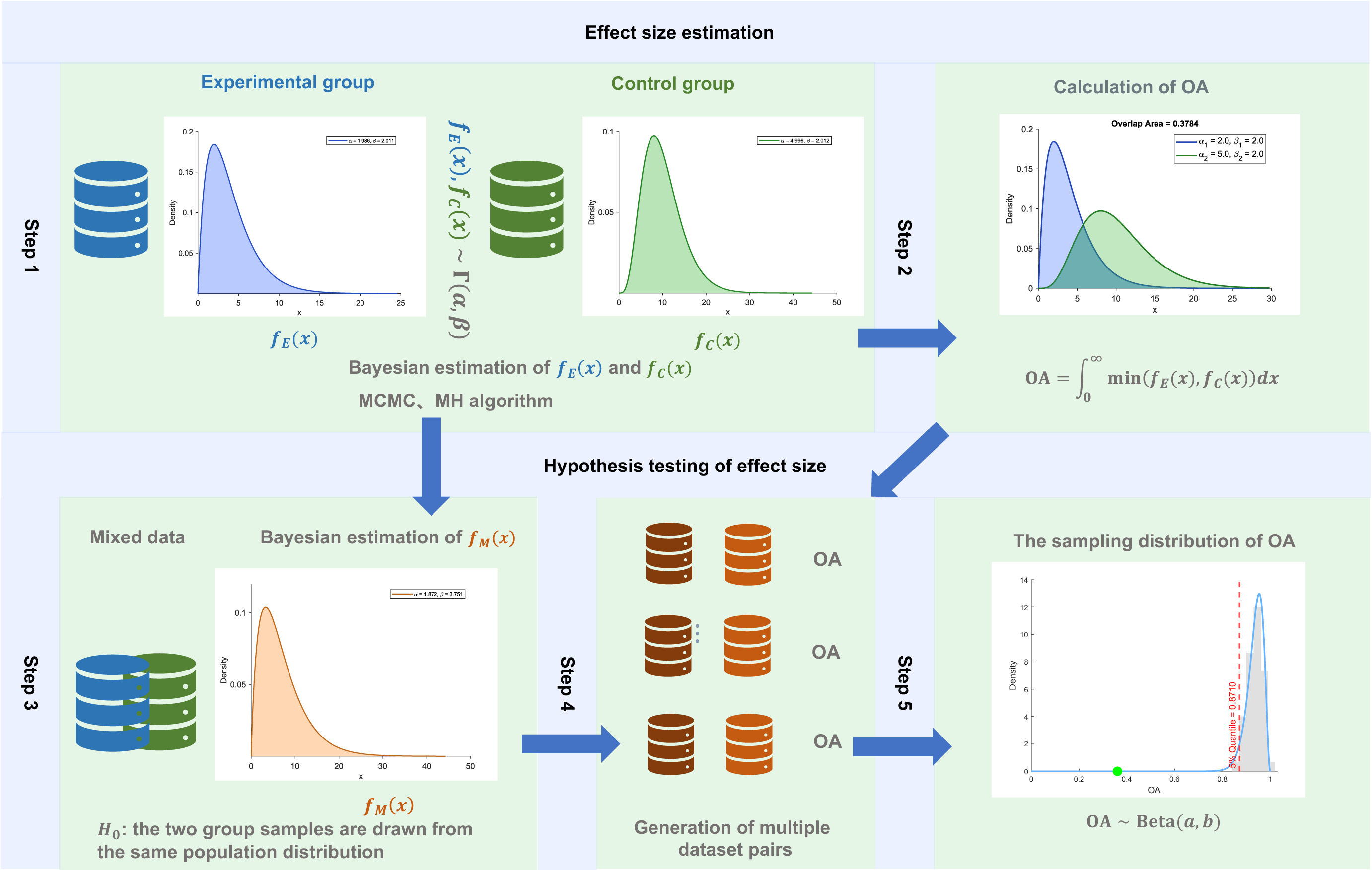
Flowchart illustrating OA estimation and hypothesis testing. *MCMC, Markov chain Monte Carlo; MH, Metropolis-Hastings.*

#### Step 1: Bayesian estimation of the population distributions for the experimental and control groups

Assuming that the dataset consists of independent and identically distributed (i.i.d.) observations, which follow a Gamma distribution. We perform Bayesian estimation to infer the population distribution for each group.

#### Step 2: Calculation of the OA to measure effect size

Based on Step 1, we calculated the OA between the population distributions of the two groups, which intuitively quantifies the degree of difference.

#### Step 3: Construction of the null distribution and Bayesian estimation

Assuming the null hypothesis that both groups are drawn from the same population, we pool the observed data to estimate the common population distribution, following Step 1.

#### Step 4: Generation of multiple dataset pairs and the OA estimation for each pair

Based on the null distribution obtained in Step 3, we generate multiple pairs of datasets from the null distribution and calculate the OA for each pair following Steps 1 and 2.

#### Step 5: Fitting the sampling distribution of the OA

Since the OA values range between 0 and 1, we assume their sampling distribution follows a Beta distribution. Based on the OA values obtained in Step 4, we fit a Beta distribution.

Based on the Beta distribution obtained in Step 5, we calculate *p*-values and determine thresholds to assess the significance of effects. For replicated experiments, we re-estimate the OA and assess its statistical significance based on the threshold.

### Sensitivity and robustness analyses

A series of simulation studies was conducted to comprehensively assess the sensitivity and robustness of the OA and to compare its performance with the traditional Cohen’s *d* approach. The simulations are designed to validate the advantages of the proposed method under various conditions. Both groups are assumed to have Gamma distributions. Figure 2-(a) illustrates Cases 1 to 4, each with different parameter settings. In Cases 1 to 3, the experimental and control groups exhibit varying degrees of distributional differences. In Case 4, the two groups are the same. The performance of estimation and hypothesis testing methods based on OA is influenced by both sample size and the number of resampling observations. Therefore, this study investigates the impact of varying sample sizes *n*_1_ = *n*_2_ = *n* = 10, 20, 30, 50,100 and NR = 20 or 50 to evaluate the performance of Bayesian parameter estimation and associated hypothesis testing.

**Figure 2.**
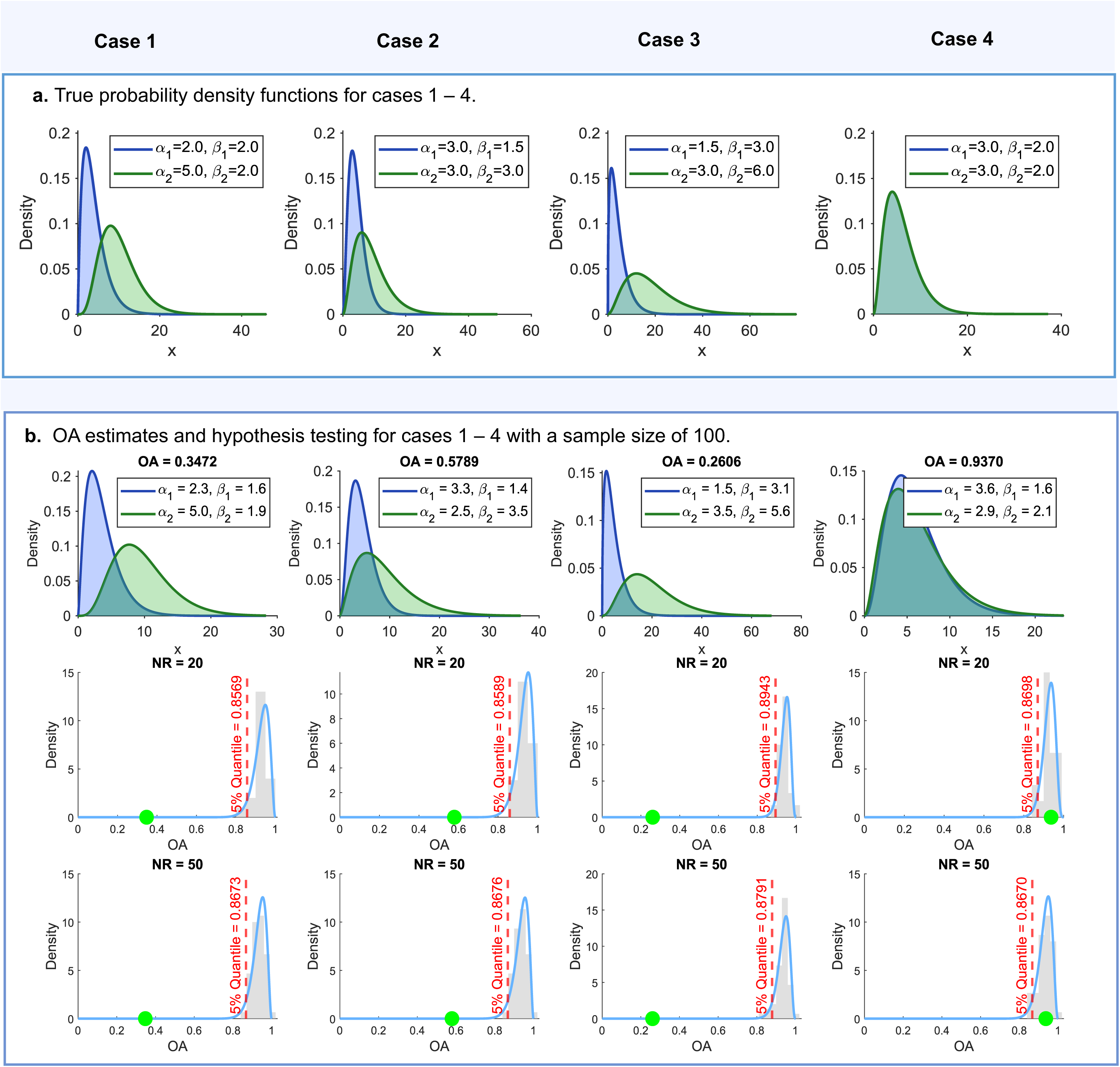
Sensitivity analysis of the OA method. **a.** The true probability density functions of Gamma distributions under four different cases. Case 1: Different shapes *⍺*, same scale *β*,***x***_1_ ∼ *Γ*(*⍺*_1_ = 2, *β*_1_ = 2), ***x***_2_ ∼ *Γ*(*⍺*_2_ = 5, *β*_2_ = 2); Case 2: Same shape *⍺*, different scales *β*, ***x***_1_ ∼ *Γ*(*⍺*_1_ = 3, *β*_1_ = 1.5), ***x***_2_ ∼ *Γ*(*⍺*_2_ = 3, *β*_2_ = 3); Case 3: Different shapes *⍺* and scales *β*, ***x***_1_ ∼ *Γ*(*⍺*_1_ = 1.5, *β*_1_ = 3), ***x***_2_ ∼ *Γ*(*⍺*_2_ = 3, *β*_2_ = 6); Case 4: Same shape *⍺* and scale *β*, ***x***_1_ ∼ *Γ*(*⍺*_1_ = 3, *β*_1_ = 2), ***x***_2_ ∼ *Γ*(*⍺*_2_ = 3, *β*_2_ = 2). **b.** The estimates of two population distributions and their OA in a single random simulation, Group 1, blue curve; Group 2, green curve; the shaded area represents the OA (top row), along with the sampling distributions of OA from NR = 20 (middle row) and NR = 50 (bottom row) based on different cases and a sample size of *n* = 100; green dots, values of the OA; red dashed line, threshold at the 5% significance level.

Taking Cases 1 and 4 from Table 1 as examples, we observe that as the sample size increases, estimates of population parameters get closer to their true values, and standard errors decrease. This means larger samples give more reliable estimates. Cohen’s *d* is much less stable with small samples, showing higher standard errors and biased estimates of the effect size. For instance, in Case 4 with a sample size of 10, Cohen’s *d* is estimated at 0.0563 with a standard error of 0.4996. According to the 3*σ* rule, such uncertainty is sufficient for a random experiment to detect a spurious effect even when no true effect exists, undermining the reliability of effect detection and the reproducibility of research findings. The OA is more robust in small samples; under the same conditions, its estimate is 0.8382 with a standard error of 0.0944, indicating that random fluctuations have a lesser impact. The *p*-values from Case 1 and Case 4 also illustrate how NR affects the results: as NR increases, the *p*-value decreases in Case 1, increasing the likelihood of rejecting the null hypothesis, whereas it increases in Case 4, increasing the likelihood of accepting the null hypothesis. This pattern indicates that larger NR enhances the stability of statistical testing, and Cases 2 and 3 show trends consistent with Case 1. Additionally, Figure 2-(b) presents estimates of the two population distributions, OA values, and the corresponding Beta distributions for hypothesis testing with a sample size of 100 across different cases. It shows that the estimated population distributions of the two groups generally align with the true distributions. In Cases 1–3, the OA values deviate substantially from the 5% significance threshold under the null hypothesis, whereas in Case 4 the OA value falls within the acceptance region, indicating that the method can effectively detect significant effects while maintaining high reliability when no effect exists. Results for other sample sizes are provided in Figure S1.

**Table 1:**
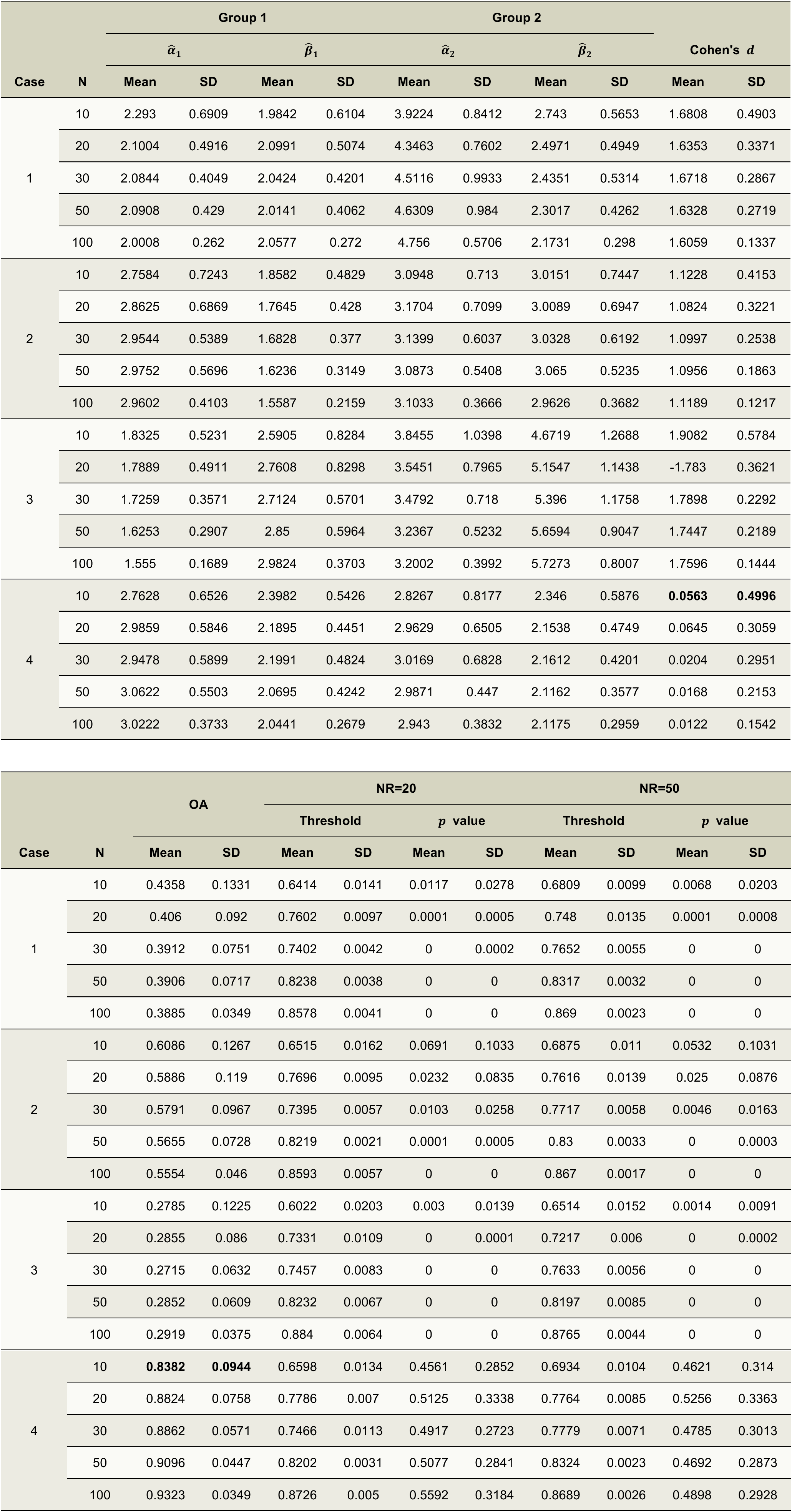
Estimates of Gamma parameters, Cohen’s *d*, OA, 5% percentile of the Beta distribution, and *p*-values across different cases and sample sizes. All results presented are the means and the standard deviations of 100 repeated simulations.

| Case | N | Group 1 | | | | Group 2 | | | | Cohen's $d$ | |
| --- | --- | --- | --- | --- | --- | --- | --- | --- | --- | --- | --- |
| | | $\hat{\alpha}_1$ | | $\hat{\beta}_1$ | | $\hat{\alpha}_2$ | | $\hat{\beta}_2$ | | Cohen's $d$ | |
|  |  | Mean | SD | Mean | SD | Mean | SD | Mean | SD | Mean | SD |
| 1 | 10 | 2.293 | 0.6909 | 1.9842 | 0.6104 | 3.9224 | 0.8412 | 2.743 | 0.5653 | 1.6808 | 0.4903 |
|  | 20 | 2.1004 | 0.4916 | 2.0991 | 0.5074 | 4.3463 | 0.7602 | 2.4971 | 0.4949 | 1.6353 | 0.3371 |
|  | 30 | 2.0844 | 0.4049 | 2.0424 | 0.4201 | 4.5116 | 0.9933 | 2.4351 | 0.5314 | 1.6718 | 0.2867 |
|  | 50 | 2.0908 | 0.429 | 2.0141 | 0.4062 | 4.6309 | 0.984 | 2.3017 | 0.4262 | 1.6328 | 0.2719 |
|  | 100 | 2.0008 | 0.262 | 2.0577 | 0.272 | 4.756 | 0.5706 | 2.1731 | 0.298 | 1.6059 | 0.1337 |
| 2 | 10 | 2.7584 | 0.7243 | 1.8582 | 0.4829 | 3.0948 | 0.713 | 3.0151 | 0.7447 | 1.1228 | 0.4153 |
|  | 20 | 2.8625 | 0.6869 | 1.7645 | 0.428 | 3.1704 | 0.7099 | 3.0089 | 0.6947 | 1.0824 | 0.3221 |
|  | 30 | 2.9544 | 0.5389 | 1.6828 | 0.377 | 3.1399 | 0.6037 | 3.0328 | 0.6192 | 1.0997 | 0.2538 |
|  | 50 | 2.9752 | 0.5696 | 1.6236 | 0.3149 | 3.0873 | 0.5408 | 3.065 | 0.5235 | 1.0956 | 0.1863 |
|  | 100 | 2.9602 | 0.4103 | 1.5587 | 0.2159 | 3.1033 | 0.3666 | 2.9626 | 0.3682 | 1.1189 | 0.1217 |
| 3 | 10 | 1.8325 | 0.5231 | 2.5905 | 0.8284 | 3.8455 | 1.0398 | 4.6719 | 1.2688 | 1.9082 | 0.5784 |
|  | 20 | 1.7889 | 0.4911 | 2.7608 | 0.8298 | 3.5451 | 0.7965 | 5.1547 | 1.1438 | -1.783 | 0.3621 |
|  | 30 | 1.7259 | 0.3571 | 2.7124 | 0.5701 | 3.4792 | 0.718 | 5.396 | 1.1758 | 1.7898 | 0.2292 |
|  | 50 | 1.6253 | 0.2907 | 2.85 | 0.5964 | 3.2367 | 0.5232 | 5.6594 | 0.9047 | 1.7447 | 0.2189 |
|  | 100 | 1.555 | 0.1689 | 2.9824 | 0.3703 | 3.2002 | 0.3992 | 5.7273 | 0.8007 | 1.7596 | 0.1444 |
| 4 | 10 | 2.7628 | 0.6526 | 2.3982 | 0.5426 | 2.8267 | 0.8177 | 2.346 | 0.5876 | **0.0563** | **0.4996** |
|  | 20 | 2.9859 | 0.5846 | 2.1895 | 0.4451 | 2.9629 | 0.6505 | 2.1538 | 0.4749 | 0.0645 | 0.3059 |
|  | 30 | 2.9478 | 0.5899 | 2.1991 | 0.4824 | 3.0169 | 0.6828 | 2.1612 | 0.4201 | 0.0204 | 0.2951 |
|  | 50 | 3.0622 | 0.5503 | 2.0695 | 0.4242 | 2.9871 | 0.447 | 2.1162 | 0.3577 | 0.0168 | 0.2153 |
|  | 100 | 3.0222 | 0.3733 | 2.0441 | 0.2679 | 2.943 | 0.3832 | 2.1175 | 0.2959 | 0.0122 | 0.1542 |

| Case | N | OA |  | NR=20 |  |  |  | NR=50 |  |  |  |
| --- | --- | --- | --- | --- | --- | --- | --- | --- | --- | --- | --- |
| | | Mean | SD | Threshold | | $p$ value | | Threshold | | $p$ value | |
|  |  |  |  | Mean | SD | Mean | SD | Mean | SD | Mean | SD |
| 1 | 10 | 0.4358 | 0.1331 | 0.6414 | 0.0141 | 0.0117 | 0.0278 | 0.6809 | 0.0099 | 0.0068 | 0.0203 |
|  | 20 | 0.406 | 0.092 | 0.7602 | 0.0097 | 0.0001 | 0.0005 | 0.748 | 0.0135 | 0.0001 | 0.0008 |
|  | 30 | 0.3912 | 0.0751 | 0.7402 | 0.0042 | 0 | 0.0002 | 0.7652 | 0.0055 | 0 | 0 |
|  | 50 | 0.3906 | 0.0717 | 0.8238 | 0.0038 | 0 | 0 | 0.8317 | 0.0032 | 0 | 0 |
|  | 100 | 0.3885 | 0.0349 | 0.8578 | 0.0041 | 0 | 0 | 0.869 | 0.0023 | 0 | 0 |
| 2 | 10 | 0.6086 | 0.1267 | 0.6515 | 0.0162 | 0.0691 | 0.1033 | 0.6875 | 0.011 | 0.0532 | 0.1031 |
|  | 20 | 0.5886 | 0.119 | 0.7696 | 0.0095 | 0.0232 | 0.0835 | 0.7616 | 0.0139 | 0.025 | 0.0876 |
|  | 30 | 0.5791 | 0.0967 | 0.7395 | 0.0057 | 0.0103 | 0.0258 | 0.7717 | 0.0058 | 0.0046 | 0.0163 |
|  | 50 | 0.5655 | 0.0728 | 0.8219 | 0.0021 | 0.0001 | 0.0005 | 0.83 | 0.0033 | 0 | 0.0003 |
|  | 100 | 0.5554 | 0.046 | 0.8593 | 0.0057 | 0 | 0 | 0.867 | 0.0017 | 0 | 0 |
| 3 | 10 | 0.2785 | 0.1225 | 0.6022 | 0.0203 | 0.003 | 0.0139 | 0.6514 | 0.0152 | 0.0014 | 0.0091 |
|  | 20 | 0.2855 | 0.086 | 0.7331 | 0.0109 | 0 | 0.0001 | 0.7217 | 0.006 | 0 | 0.0002 |
|  | 30 | 0.2715 | 0.0632 | 0.7457 | 0.0083 | 0 | 0 | 0.7633 | 0.0056 | 0 | 0 |
|  | 50 | 0.2852 | 0.0609 | 0.8232 | 0.0067 | 0 | 0 | 0.8197 | 0.0085 | 0 | 0 |
|  | 100 | 0.2919 | 0.0375 | 0.884 | 0.0064 | 0 | 0 | 0.8765 | 0.0044 | 0 | 0 |
| 4 | 10 | **0.8382** | **0.0944** | 0.6598 | 0.0134 | 0.4561 | 0.2852 | 0.6934 | 0.0104 | 0.4621 | 0.314 |
|  | 20 | 0.8824 | 0.0758 | 0.7786 | 0.007 | 0.5125 | 0.3338 | 0.7764 | 0.0085 | 0.5256 | 0.3363 |
|  | 30 | 0.8862 | 0.0571 | 0.7466 | 0.0113 | 0.4917 | 0.2723 | 0.7779 | 0.0071 | 0.4785 | 0.3013 |
|  | 50 | 0.9096 | 0.0447 | 0.8202 | 0.0031 | 0.5077 | 0.2841 | 0.8324 | 0.0023 | 0.4692 | 0.2873 |
|  | 100 | 0.9323 | 0.0349 | 0.8726 | 0.005 | 0.5592 | 0.3184 | 0.8689 | 0.0026 | 0.4898 | 0.2928 |

Figure 3 further illustrates the advantages of the OA method and its sensitivity to changes in sample size. As shown in Figure 3-(a,b), with increasing sample size, the estimates from 100 simulations cover the true values, and the widths of the OA boxplots decrease, indicating greater stability in the estimates. However, when the sample size is below 30, the parameter estimates for Group 2 in Case 1 and Case 3 show slightly poorer performance. The larger variance in the population distributions and the pronounced differences in the shape parameter *⍺* and rate parameter *β* between the two groups make the estimates more susceptible to small-sample fluctuations. Although the parameter estimates are somewhat unstable under small sample sizes, the hypothesis testing framework based on the OA method can largely compensate for this instability. As shown in Figure 3-(c), even with a sample size of *n* = 10, the *p*-values from 100 repeated experiments in Case 1 and Case 3 are consistently below 0.05, indicating that the test results are not significantly affected by the small sample size. By contrast, at *n* = 10, because the differences in *⍺* and *β* between the two groups are smaller in Case 2, the corresponding *p*-values are not consistently below 0.05. Notably, increasing the NR value leads to narrower *p*-value boxplots, enhancing the stability and reliability of the hypothesis testing results. Accordingly, our method enables rigorous inference on the statistical significance of effect sizes, addressing a limitation of Cohen’s *d*.

**Figure 3.**
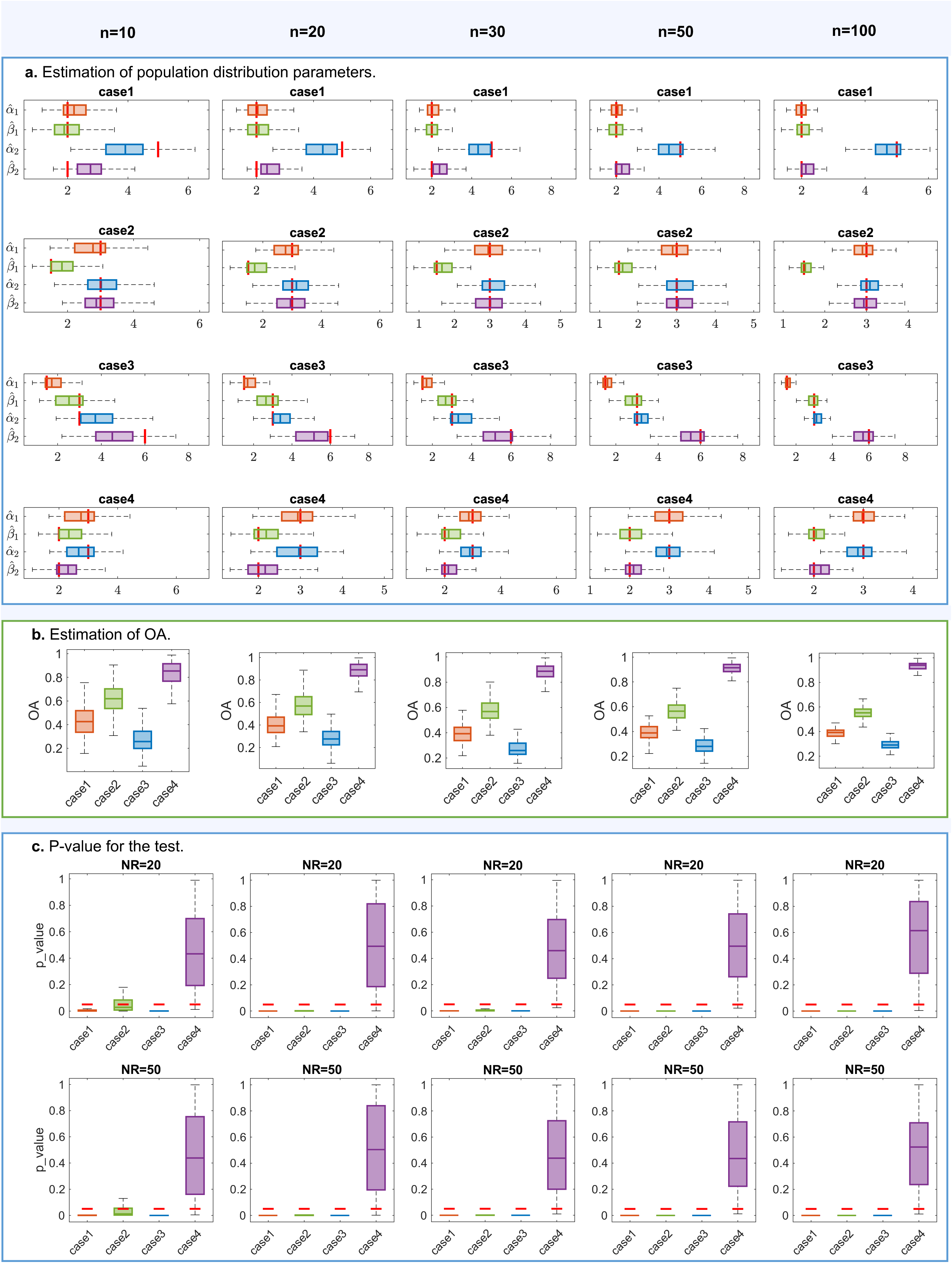
Boxplots of estimated two population parameters, OA values, and hypothesis-test results across cases and sample sizes from 100 random simulations. **a.** Boxplots of the estimated Gamma distribution parameters for the Experimental and Control groups. Red lines indicate the true parameter values. **b.** Boxplots of OA values. **c.** Boxplots of *p*- values under two settings (NR *=* 20 and NR *=* 50). Red horizontal lines at *p* = 0.05 indicate the significance threshold.

From the preceding results, it is evident that in Cases 1– 4 the effect size estimated by Cohen’s *d* is susceptible to random fluctuations under small sample sizes, leading to biased estimates. As the sample size increases, however, the estimates become more stable and better reflect the true effect. Therefore, it is necessary to further examine scenarios in which Cohen’s *d* may still perform poorly even under large sample conditions. We build on the setup of Case 1 by first fixing the distribution of the first group as ***x***_1_ ∼ Γ(*⍺*_1_ = 2, *β*_1_ = 2). Under the constraint that Cohen’s *d* equals zero, that is, in situations where the method is unable to detect any effect, we systematically explore all possible parameter combinations for the second group (as shown in Figure 4-(a)) to identify the parameter region in which Cohen’s *d* breaks down despite large samples. Based on this exploration, we select the second group to follow ***x***_2_ ∼ Γ(*⍺*_2_ = 8, *β*_2_ = 0.5), which lies within the identified failure region. Using this setup, we then compare the performance of Cohen’s *d* and the OA method under both balanced (Case 5: *n* = 100) and unbalanced (Case 6: *n*_1_ = 100 vs. *n*_2_ = 50) sample sizes, allowing us to evaluate the advantages of the OA method in contexts where Cohen’s *d* is ineffective.

**Figure 4.**
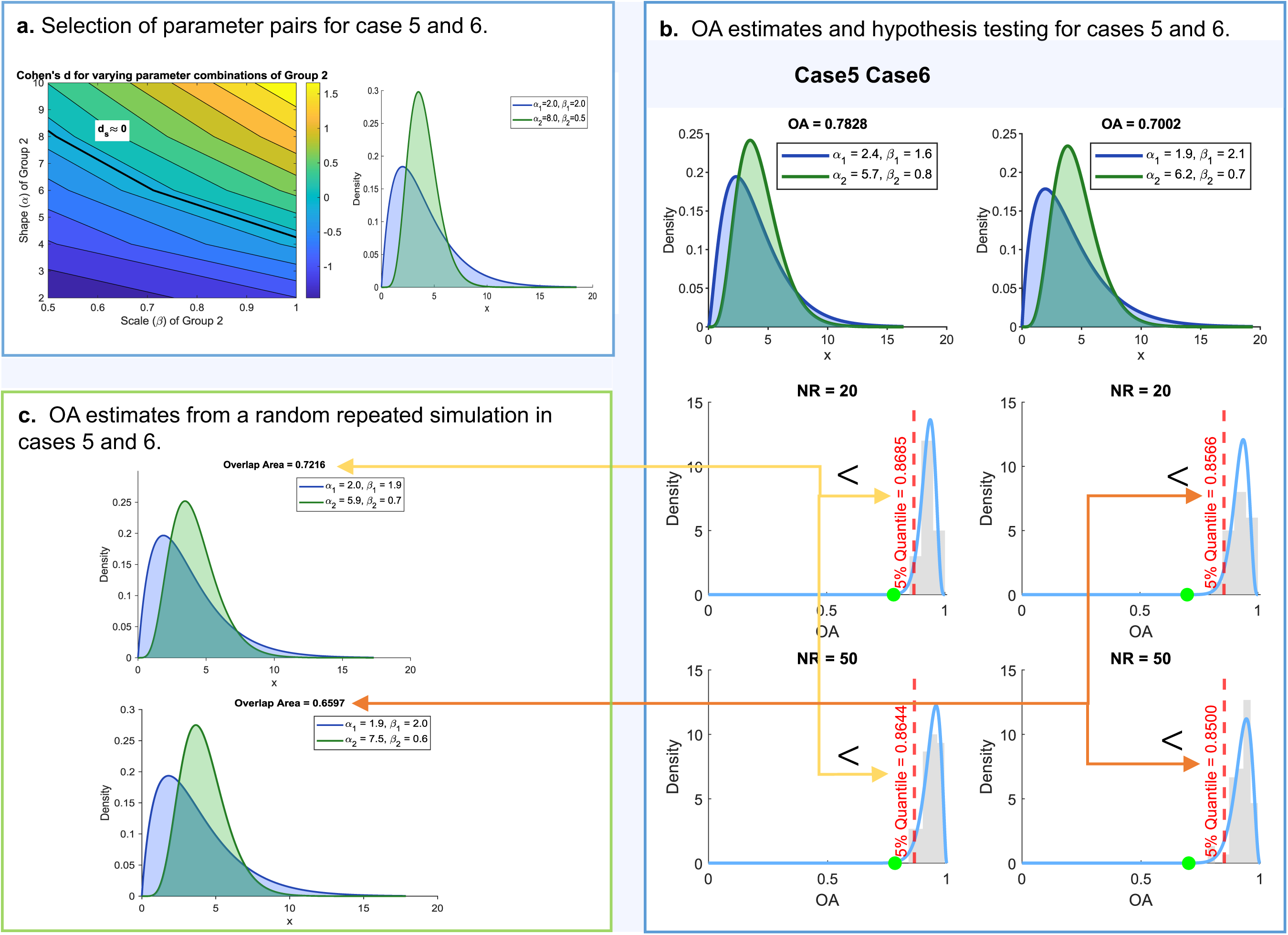
Robustness analysis of the OA method. **a.** Parameter pairs with Cohen’s *d* = 0 (left); probability density functions of the Gamma distributions ***x***_1_ ∼ *Γ*(*⍺*_1_ = 2, *β*_1_ = 2), ***x***_2_ ∼ *Γ*(*⍺*_2_ = 8, *β*_2_ = 0.5) under Cases 5 and 6 (right). **b.** OA estimate from a single random simulation with balanced (Case 5: *n*_1_=*n*_2_=100) and unbalanced (Case 6: *n*_1_= 100 vs. *n*_2_ = 50) sample sizes, along with the corresponding hypothesis-test results. **c.** The re-estimates of OA from a random replication, compared with the threshold values under the null hypothesis. The yellow arrow denotes the comparison between the re-estimate of OA and the specified threshold value under NR *=* 20 and NR *=* 50.

As shown in Table 2, Cohen’s *d* is only 0.0205 in Case 5 and even decreases to −0.0028 in Case 6, indicating that it completely fails to capture the effect size, ‘masking’ the true effect. By contrast, the OA method demonstrates strong stability and effectiveness in estimating the population parameters, the OA values, and in conducting hypothesis testing; the resulting *p*-values are far below 0.05. This indicates that, whether the sample sizes are balanced or unbalanced, the OA method reliably quantifies effect sizes and provides valid significance testing.

**Table 2:** Estimates of Gamma parameters, Cohen’s *d*, OA, 5% percentile of the Beta distribution, and *p*-values across different cases and sample sizes. All results presented are the mean and the standard deviations of 100 repeated simulations.

|  |  | Group 1 |  | Group 2 |  |  |  | NR = 20 |  | NR = 50 |  |
| --- | --- | --- | --- | --- | --- | --- | --- | --- | --- | --- | --- |
| | Case | $\hat{\alpha}_1$ | $\hat{\beta}_1$ | $\hat{\alpha}_2$ | $\hat{\beta}_2$ | Cohen's $d$ | OA | Threshold | $p$ value | Threshold | $p$ value |
| Mean | 5 | 2.0261 | 2.033 | 7.1509 | 0.5852 | **0.0205** | 0.6868 | 0.8706 | **0.0000** | 0.8665 | **0.0001** |
|  | 6 | 2.0599 | 2.0015 | 6.3399 | 0.6733 | **-0.0028** | 0.7178 | 0.8568 | **0.0008** | 0.8482 | **0.0017** |
| SD | 5 | 0.2481 | 0.2972 | 1.0027 | 0.0828 | 0.1460 | 0.0410 | 0.0032 | 0.0001 | 0.0015 | 0.0003 |
|  | 6 | 0.2443 | 0.2570 | 1.0736 | 0.1143 | 0.1427 | 0.0463 | 0.0023 | 0.0058 | 0.0041 | 0.0090 |

To further validate that our method can reliably detect significance in replication studies, we use the threshold derived from the OA sampling distribution of Case 5 and 6. We then generate a new random dataset to evaluate the method’s ability to identify significant differences in replication experiments. As shown in Figure 4-(b,c), OA = 0.7216, OA = 0.6597 obtained from a new random simulation dataset is significantly lower than the 5% significance thresholds based on Case 5 ( 0.8685 for NR = 20 and 0.8644 for NR = 50) and 6 (0.8566 for NR = 20 and 0.8500 for NR = 50). This result demonstrates that the OA method can consistently detect significant effects in replication experiments.

### The effect of environmental enrichment on the mouse brain synaptome

We applied OA measurement to a synaptome mapping dataset generated from mice raised in environmental enrichment (EE) or standard caging (SC) from postnatal day 0 to 90^16,17^. Environmental enrichment is a well-established paradigm in neuroscience incorporating enhanced sensory, cognitive, social, and physical stimulation. It has been shown to alter synaptic and neuronal properties across brain regions, but prior studies have lacked the resolution to quantify experience-dependent plasticity at single-synapse scale. This dataset—comprising brainwide, high-resolution profiles of excitatory synapse types—was generated using the synaptome mapping approach for PSD95^18^. Our primary aim was to determine whether we could detect changes in the mouse brain synaptome (as reflected in synaptic density, intensity, or size) driven by EE and to assess the regionality and scale of any such effects.

Based on these observations, we applied the aforementioned analytical approach to comprehensively evaluate the distribution patterns of synapse density, intensity, and size across each brain subregion. Through effect size estimation and hypothesis testing, we aimed to identify brain regions that exhibit significant changes in response to environmental influences. Furthermore, by integrating the population distribution maps of both groups, we explored potential causes and mechanisms underlying these differences.

Figure 5-(a) shows the distributions of Cohen’s *d* and the OA between the two groups, demonstrating that our proposed method can effectively identify brain regions exhibiting significant differences between the two environmental conditions. Notably, our method shows a clear advantage in detecting differences in PSD95 intensity, where Cohen’s *d* alone fails to reveal such distinctions. Across multiple brain regions, PSD95 intensity—quantified as the mean pixel intensity of PSD95-labelled synapses—differed markedly between EE and SC groups. These intensity values reflect the average concentration of PSD95 protein per synapse within each region and can be interpreted as a proxy for the molecular composition of the postsynaptic terminal. PSD95 is a core scaffolding protein in excitatory synapses that anchors and regulates neurotransmitter receptors and associated signalling complexes^19,20^. Prior studies have demonstrated that environmental enrichment can modulate PSD95 expression at the mRNA and protein levels across cortical, hippocampal, and striatal regions ^21–23^. High-resolution imaging studies have further shown that EE can reduce variability in PSD95 punctum size and increase spine head area, suggesting more homogeneous synaptic populations under enriched conditions^24^. In this context, regional differences in PSD95 intensity may reflect experience-dependent molecular reorganisation of the synaptome, capturing subtle functional adaptations that are not detectable through changes in synapse number or size alone.

**Figure 5.**
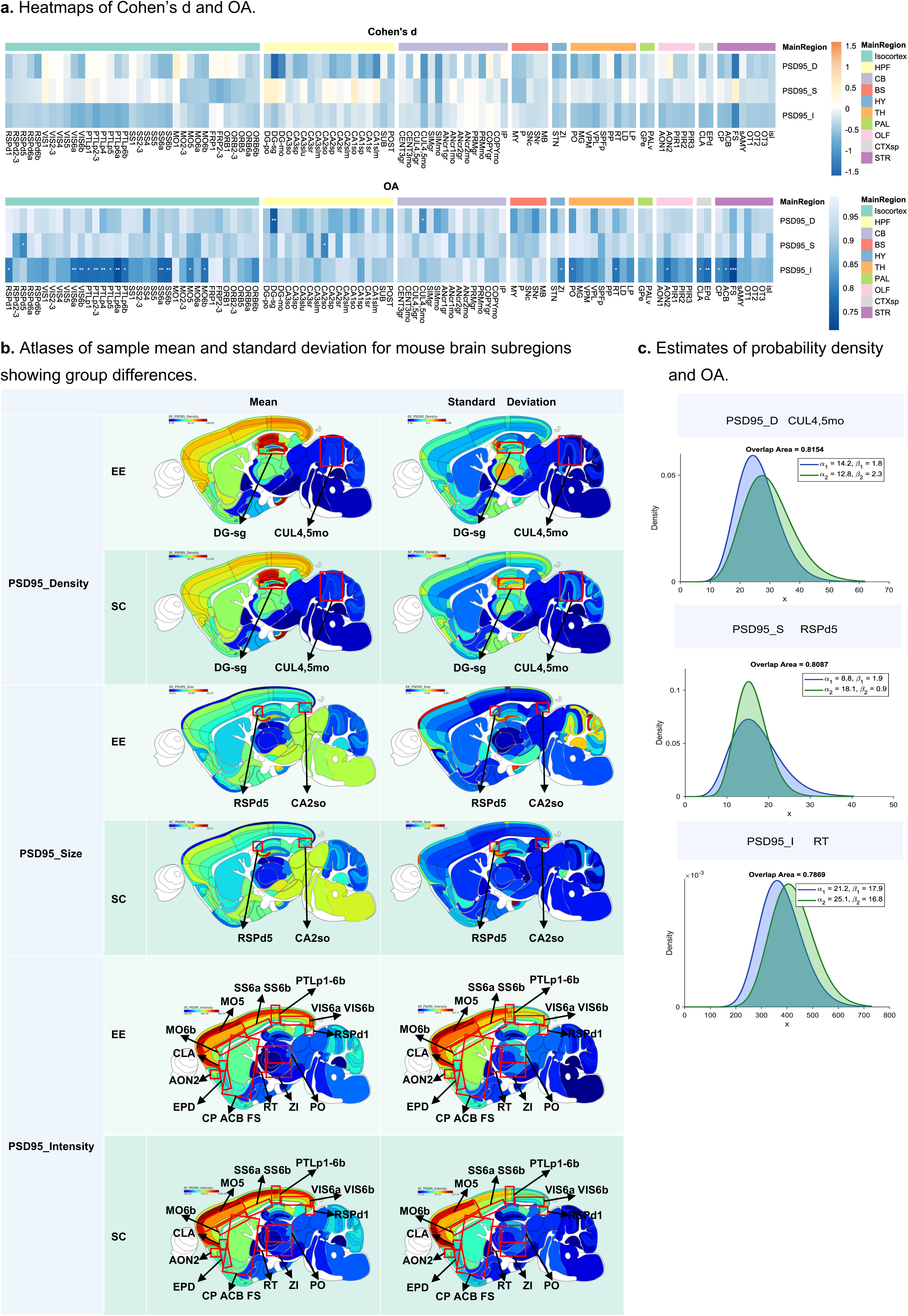
Analysis of the effect of environmental enrichment on the mouse brain synaptome. **a.** Heatmap of Cohen’s *d* (upper panel) and the OA (lower panel) between the EE and SC groups across 101 brain subregions for three synaptic measures, including density (D), size (S), and intensity (I) of PSD95. Asterisks indicate statistically significant differences between the two groups at the corresponding brain subregion: ∗ *p* < 0.1, ∗∗ *p* < 0.05, *** *p* < 0.01. **b.** Atlases of the mean and standard deviation of PSD95 density, size, and intensity. **c.** Three examples of probability density distributions showing differences. Top to bottom: Both the mean and variance show differences (PSD95 density in CUL4,5mo); Differences in variance(PSD95 size in RSPd5); Differences in mean (PSD95 intensity in RT).

Figure 5-(b) displays sample mean and standard deviation maps of synaptic density, size, and intensity for PSD95 across 101 brain subregions under the different environmental conditions. Although the mean maps show no visually obvious group differences between EE and SC, the standard deviation maps reveal notable differences in the variability of synaptic features, highlighting the impact of environmental conditions on the mouse brain synaptome.

In Figure 5-(c), we present representative examples of three distinct types of distributional differences among brain regions that exhibited significant changes; detailed distribution difference plots for the remaining brain regions are provided in Figure S2. The first type is represented by PSD95 density in the CUL4,5mo brain region, where the EE group shows both a lower mean and reduced variance compared with the SC group. The second type is represented by PSD95 size in the RSPd5 region, where the mean values between the EE and SC groups are similar, but the variance is markedly higher in the EE group, suggesting that while the average size of PSD95-expressing synapses remains unchanged, individual differences within the EE group are more pronounced. The third type is represented by PSD95 intensity in the RT region, where the difference is mainly driven by the mean: the EE group exhibits a lower average intensity than the SC group, with no significant difference in variance. Together, these three types of differences highlight the diverse ways in which environmental conditions can influence synaptic organization and plasticity— either by shifting overall expression levels, modulating individual variability, or both—depending on the specific functional and anatomical properties of the brain region involved.

### OAT: an open source application toolbox

To enhance the practicality of the OA method and facilitate researchers’ efficient use across different research fields, we developed a graphical user interface (GUI) using the App Designer environment in MATLAB. The application is implemented as a .mlapp file and requires MATLAB to be installed on the user’s local machine. After launching the OA toolbox (OAT) App, an interactive window appears, as illustrated in Figure 6.

**Figure 6.**
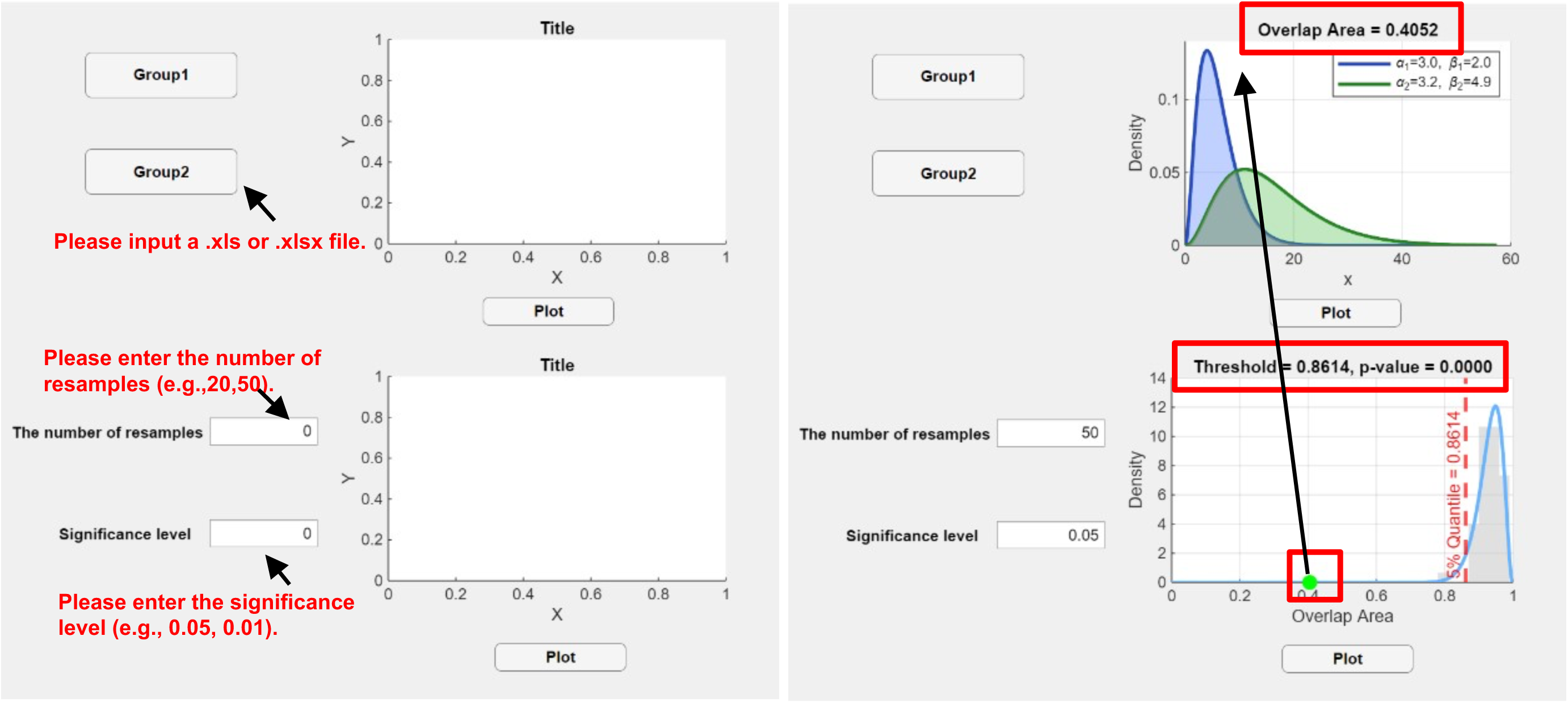
OAT user interface.

OAT accepts input data in Excel format (.xls or .xlsx). Users should prepare two datasets corresponding to Group 1 and Group 2 and save them as separate Excel files. Data import is achieved by clicking the Group1 and Group2 buttons. Upon successful data upload, OAT displays the message ‘Data upload was successful’; if no file is selected or the import fails, the message ‘No file has been selected’ will appear. Once the data are imported, users can click the Plot button at the top right of the interface to perform. Next, the user needs to enter the value of NR in the Resampling Replicates section, and specify the desired significance level. Then, click the Plot button beneath to generate the sampling distribution of the OA. To test the result of a replication, first click the Group1 and Group2 buttons to import new data, then click the Plot button above to generate the figure. After the plot is generated, check the OA value in the title and compare it with the Threshold shown in the lower plot. If the OA is smaller than the Threshold, the repeated experiment result is considered significant; otherwise, it is not. The supplementary materials include the installation package and test data.

## Methods

### Effect size – OA

Effect size can assess whether an experimental manipulation has a biologically meaningful effect. Taking two groups as an example, Group 1 receives a manipulation while Group 2 serves as the control. The effect size is used to quantify the magnitude of the manipulation’s effect. There are many effect size measures available to quantify the size of effects, among which Cohen’s *d* is one of the most commonly used. It measures the mean difference *µ*1 – *µ*2 between two populations, normalized by standard deviation *σ*, and is defined as:

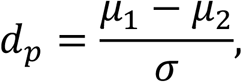

Since the population parameters *μ*_1_, *μ*_2_, and *σ* are unknown, Cohen’s *d* is estimated using sample datasets:

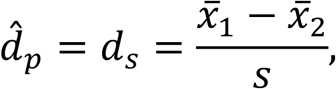

where the pooled standard deviation *s* is calculated as:

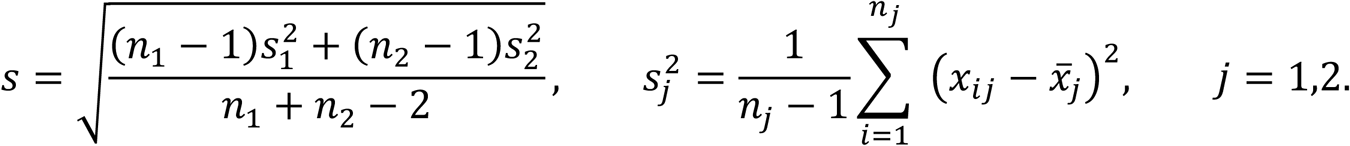

Cohen’s *d* presents several potential limitations arising from its definition. First, Cohen’s *d* relies on the assumption that the two groups have equal population variances *σ* and only reflects the difference in population means *μ*_1_ − *μ*_2_, overlooking the heterogeneity of variance between populations. When the variances are large, the true effect size may be masked. Second, the estimation of Cohen’s *d* depends on the calculation of the pooled sample variance *s*, which is influenced by the sample sizes *n*_1_, *n*_2_. When the sample sizes are small or imbalanced *n*_1_ ≠ *n*_2_, it may lead to increased bias in the estimates of the effect size. Third, given that the estimation is based on a single experimental sample dataset, the results may be influenced by random errors, reducing the likelihood of reproducing the effect size and ultimately weakening the reliability and generalizability of the findings.

Having analysed the limitations of Cohen’s *d*, we consider measuring the differences in each group’s population distributions to provide a more comprehensive representation of effect size. The assumption of the population distribution depends on the structure and characteristics of the data. Zhu and colleagues^18^ demonstrated in their synaptome mapping study that the fluorescence density, size, and intensity of the synaptic protein PSD95 can reveal the architecture and diversity of synapses throughout the mouse brain. After imaging mouse brain tissue sections, the fluorescently labelled PSD95 protein appears as a series of dispersed punctate structures (synaptic puncta) in the microscopy images. By identifying and classifying these puncta, the number of synaptic puncta per unit area is calculated as synapse density; the area of each punctum is defined as its size; and the fluorescence signal intensity of each punctum is referred to as its intensity. PSD95 consistently exhibits positive values in terms of density, size, and intensity measures. Based on this, it is reasonable to assume that the population distribution of the observed synaptic density, size, and intensity data follows a Gamma distribution^25^. The probability density function of the Gamma distribution is given by:

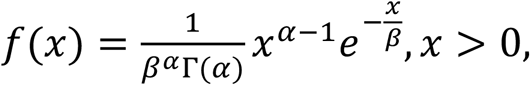

where *⍺* > 0 is the shape parameter, *β* > 0 is the scale parameter, and Γ(⋅) is the Gamma function. The notation *x* ∼ Γ(*⍺*, *β*) is used to indicate that *x* follows a Gamma distribution^26^. The skewness and concentration of a Gamma distribution can be calculated from its shape *⍺* and scale *β* parameters..

Building on this, the OA is defined between the two population distributions as follows:

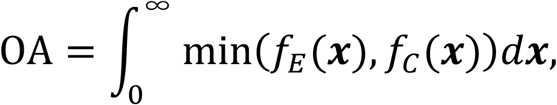

where *f*_*E*_(***x***) and *f*_*C*_(***x***) represent the Gamma distributions of the experimental and control groups, respectively. Since the integral generally has no closed form solution, the OA is calculated numerically ^27,28^.

### Bayesian estimation of population distributions

To calculate the OA, it is necessary to estimate the parameters (*⍺*_1_, *β*_1_) and (*⍺*_2_, *β*_2_) of the population distributions *f*_*E*_(***x***) and *f*_*C*_(***x***).

Let ***x*** = (*x*_1_, *x*_2_, …, *x*_*n*_) denote the observed data in a group. Assume that *x*_1_, *x*_2_, …, *x*_*n*_ are independent and identically distributed (i.i.d.) according to a Gamma distribution with shape parameter *⍺* and rate parameter *β*, i.e.

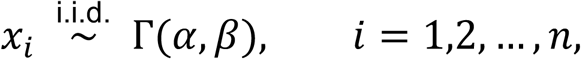

where *⍺*, *β* > 0 are unknown parameters to be estimated. Bayesian approaches are particularly useful for small-sample datasets, as they enable robust estimation by incorporating prior information. Based on Bayes’ theorem^29^, the joint posterior distribution of *⍺* and *β* given the data ***x*** is then expressed as

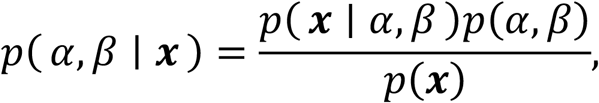

where *p*( ***x*** ∣ *⍺*, *β*) is the likelihood function, denoted as *L*(*⍺*, *β*); the prior distribution is *p*(*⍺*, *β*) = *p*(*⍺*)*p*(*β*); and *p*(***x***) is the marginal likelihood. We specify the following prior distributions for parameters:

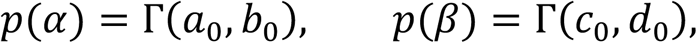

where *a*_0_, *b*_0_, *c*_0_, and *d*_0_ are given hyperparameters. Here, we assign a common value of 1 to each of them, implying weakly informative priors. Given the priors and the likelihood, the log-posterior distribution is given by:

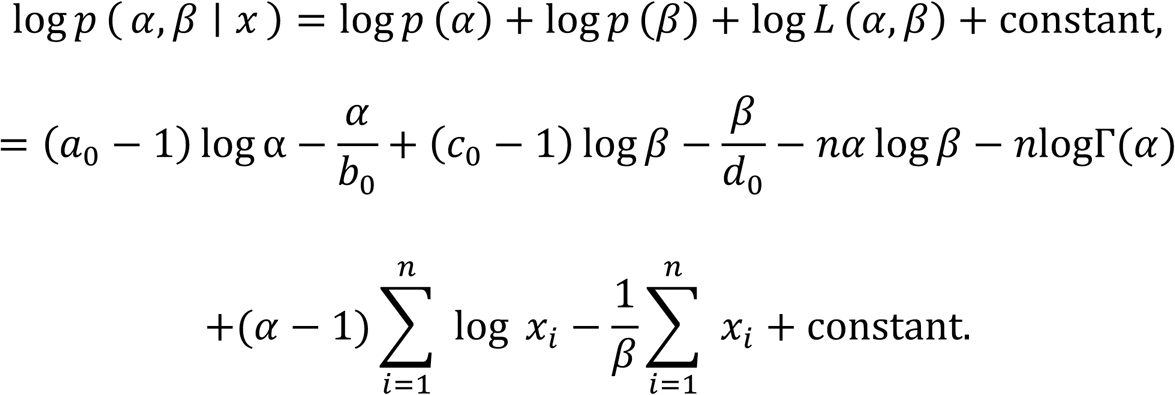

Since the posterior distribution is not analytically tractable, we employ Markov chain Monte Carlo (MCMC) ^30^ combined with the Metropolis-Hastings (MH) ^31^ algorithm to draw samples to obtain *⍺̂* and *β̂*. We estimated the distribution parameters of *f*_*E*_(***x***) and *f*_*C*_(***x***) based on the observed data from the experimental and control groups, and subsequently calculated the OA.

Using the OA as an effect size measure offers several advantages. First, it is intuitive and interpretable, directly reflecting the degree of differences between the two populations: a smaller OA indicates greater differences, whereas a larger OA suggests higher similarity. Second, it is sensitive to distributional characteristics beyond the mean and variance, capturing differences in shape such as skewness and kurtosis, providing a more comprehensive representation of population features. Third, as OA is bounded between 0 and 1, it provides an interpretable statistic and facilitates hypothesis testing through the construction of its sampling distribution.

### Hypothesis testing of OA

Assessing the significance of the OA in single experiments and replications remains a critical challenge. To address this issue, we formulate the following null hypothesis:

*H*_0_: the two group samples are drawn from the same population distribution

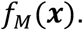

Under the null hypothesis *H*_0_, we developed a hypothesis testing framework based on the OA. Since the OA is a continuous random variable defined on [0,1], its sampling distribution can be approximated by a Beta distribution^32^, with the probability density function given by:

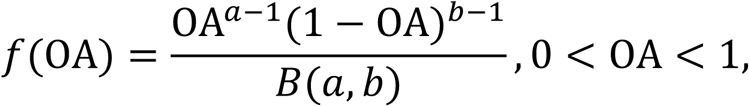

where *B*(*a*, *b*) is the Beta function with *a* and *b* as its parameters, and the OA follows a Beta(*a*, *b*) distribution. However, fitting a Beta distribution requires a series of observed OA values, i.e. OA_1_, OA_2_, …, OA_NR_, where NR denotes the number of observations. To obtain these OA values, the observed data from the experimental and control groups are pooled, and the common population distribution *f*_*M*_(***x***) is estimated. Based on the estimated distribution *f*_*M*_(***x***), we generate NR pairs of datasets through a resampling procedure^33^. As a result, we established the sampling distribution of the OA under *H*_0_ to compute *p* -value and critical thresholds, providing statistical support for significance testing in replication experiments.

## Discussion

In this study, we addressed several methodological limitations associated with conventional effect determination strategies in neuroscience by introducing a novel effect size measure, OA, together with its Bayesian estimation and hypothesis testing framework. Traditional approaches, such as Cohen’s *d*, t- tests, and Bayes factor, have provided foundational tools for quantifying and validating group differences. However, as our analysis shows, these methods often fall short in capturing complex distributional features beyond mean differences, particularly under conditions of small sample sizes, variance heterogeneity, and unbalanced group designs. These limitations not only reduce sensitivity to biologically meaningful effects but also complicate the establishment of robust significance testing criteria for replication studies.

By contrast, our proposed method enables a more nuanced assessment of effects by incorporating both mean and variance differences into a unified framework. Through simulation studies, we demonstrated that our method outperforms conventional approaches in detecting true effects under challenging conditions. When applied to a real-world brainwide dataset comparing synaptic features between mice raised in an enriched environment and standard housing, our method identified key regions with distinct effect patterns—some driven by mean shifts, others by variance changes, or both— highlighting the ability of OA to detect a broader range of biologically relevant changes in synapse populations.

Furthermore, the method supports the establishment of replicable significance thresholds, making it especially valuable for guiding follow-up experiments and increasing scientific reproducibility. To facilitate this integration, we have developed a user-friendly toolbox, OAT, which we hope will encourage wider adoption of Bayesian methods across the neuroscience community.

## Supporting information

Group1 data

Group2 data

Read_me for App

App

## Code availability

Computer code was written in MATLAB R2024b.

## Data availability

All data are available at https://www.ebi.ac.uk/biostudies/studies/S-BSST2147.

## Acknowledgements

This work was supported by a grant from the China Scholarship Council (202407030025 to Y.W.) and the National Natural Science Foundation of China (12201550 to Y.W.) and by the Wellcome Trust (302077/Z/23/Z, 218293/Z/19/Z, 221295/Z/20/Z) and Simons Initiative for the Developing Brain (SIDB) under the Simons Foundation for Autism Research Initiative (529085). For the purpose of open access, the author has applied a CC-BY public copyright licence to any Author Accepted Manuscript version arising from this submission.

## Author contributions

Y.W.: conceptualisation, methodology, formal analysis, software, visualisation, writing (original draft) and writing (review and editing). H.W.: formal analysis, investigation, resources, writing (original draft). K.H.Y, D.D., and Z.Q.: resources, visualisation, and writing (review and editing). S.G.N.G.: conceptualisation, methodology, supervision and writing (review and editing).

## Additional information

Supplemental materials have been appended to this article. The file includes (Supplementary Figure S1-S2) and App.

