## Supplemental Fig 1, 2 for "The Overlap Area as a Novel Measure of Effect Size in Neuroscience Research"

## n=10

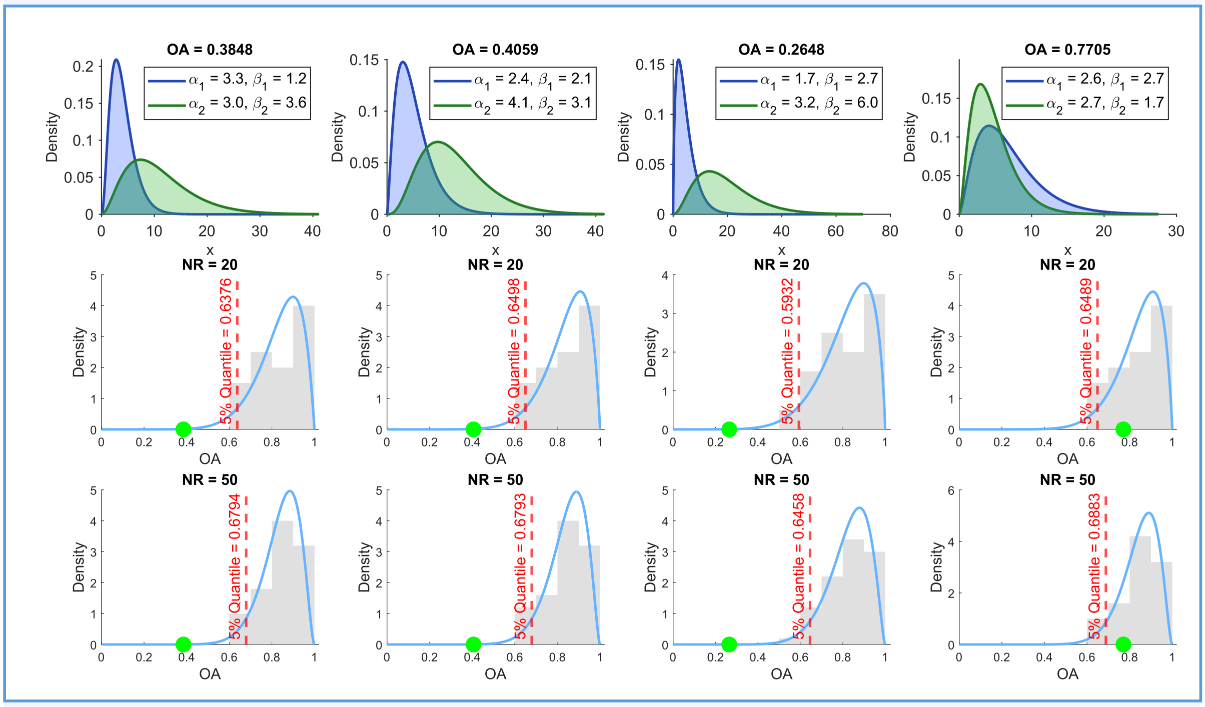

OA = 0.3848
$\alpha_1$ = 3.3, $\beta_1$ = 1.2
$\alpha_2$ = 3.0, $\beta_2$ = 3.6
OA = 0.4059
$\alpha_1$ = 2.4, $\beta_1$ = 2.1
$\alpha_2$ = 4.1, $\beta_2$ = 3.1
OA = 0.2648
$\alpha_1$ = 1.7, $\beta_1$ = 2.7
$\alpha_2$ = 3.2, $\beta_2$ = 6.0
OA = 0.7705
$\alpha_1$ = 2.6, $\beta_1$ = 2.7
$\alpha_2$ = 2.7, $\beta_2$ = 1.7
Density
x
NR = 20
5% Quantile = 0.6376
5% Quantile = 0.6498
5% Quantile = 0.5932
5% Quantile = 0.6489
NR = 50
5% Quantile = 0.6794
5% Quantile = 0.6793
5% Quantile = 0.6458
5% Quantile = 0.6883
OA


## n=20

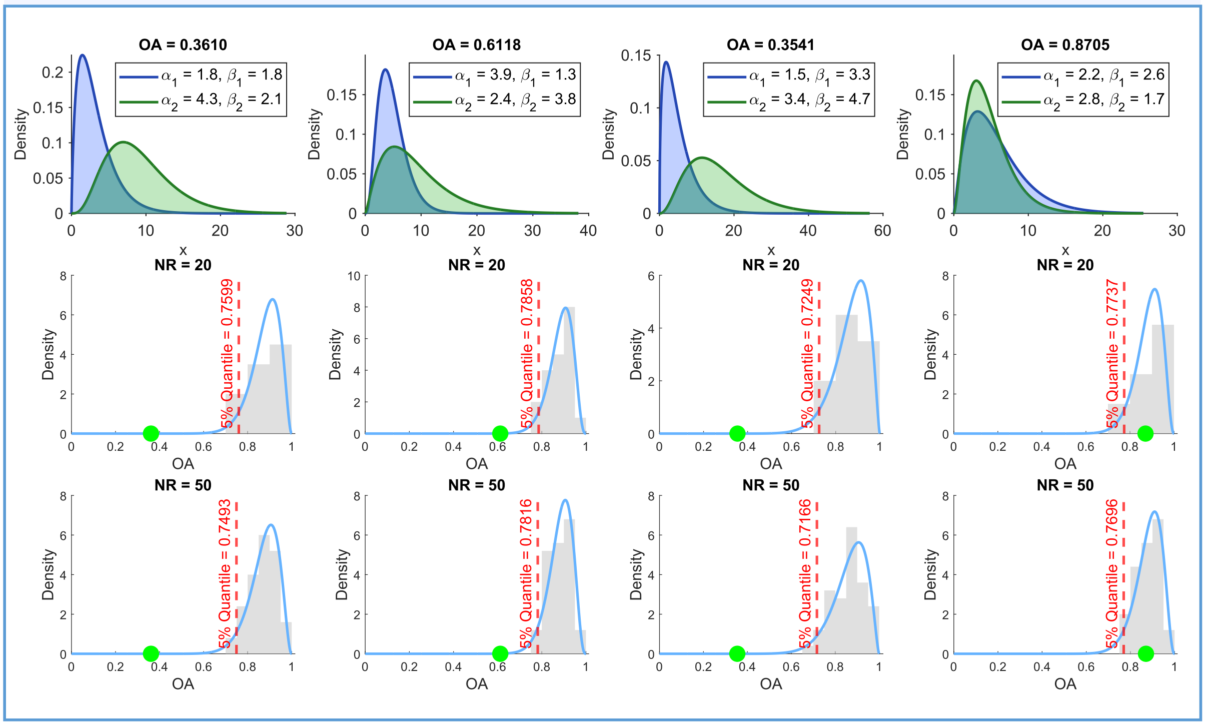

OA = 0.3610
$\alpha_1$ = 1.8, $\beta_1$ = 1.8
$\alpha_2$ = 4.3, $\beta_2$ = 2.1
OA = 0.6118
$\alpha_1$ = 3.9, $\beta_1$ = 1.3
$\alpha_2$ = 2.4, $\beta_2$ = 3.8
OA = 0.3541
$\alpha_1$ = 1.5, $\beta_1$ = 3.3
$\alpha_2$ = 3.4, $\beta_2$ = 4.7
OA = 0.8705
$\alpha_1$ = 2.2, $\beta_1$ = 2.6
$\alpha_2$ = 2.8, $\beta_2$ = 1.7
Density
x
NR = 20
5% Quantile = 0.7599
5% Quantile = 0.7858
5% Quantile = 0.7249
5% Quantile = 0.7737
NR = 50
5% Quantile = 0.7493
5% Quantile = 0.7816
5% Quantile = 0.7166
5% Quantile = 0.7696
OA

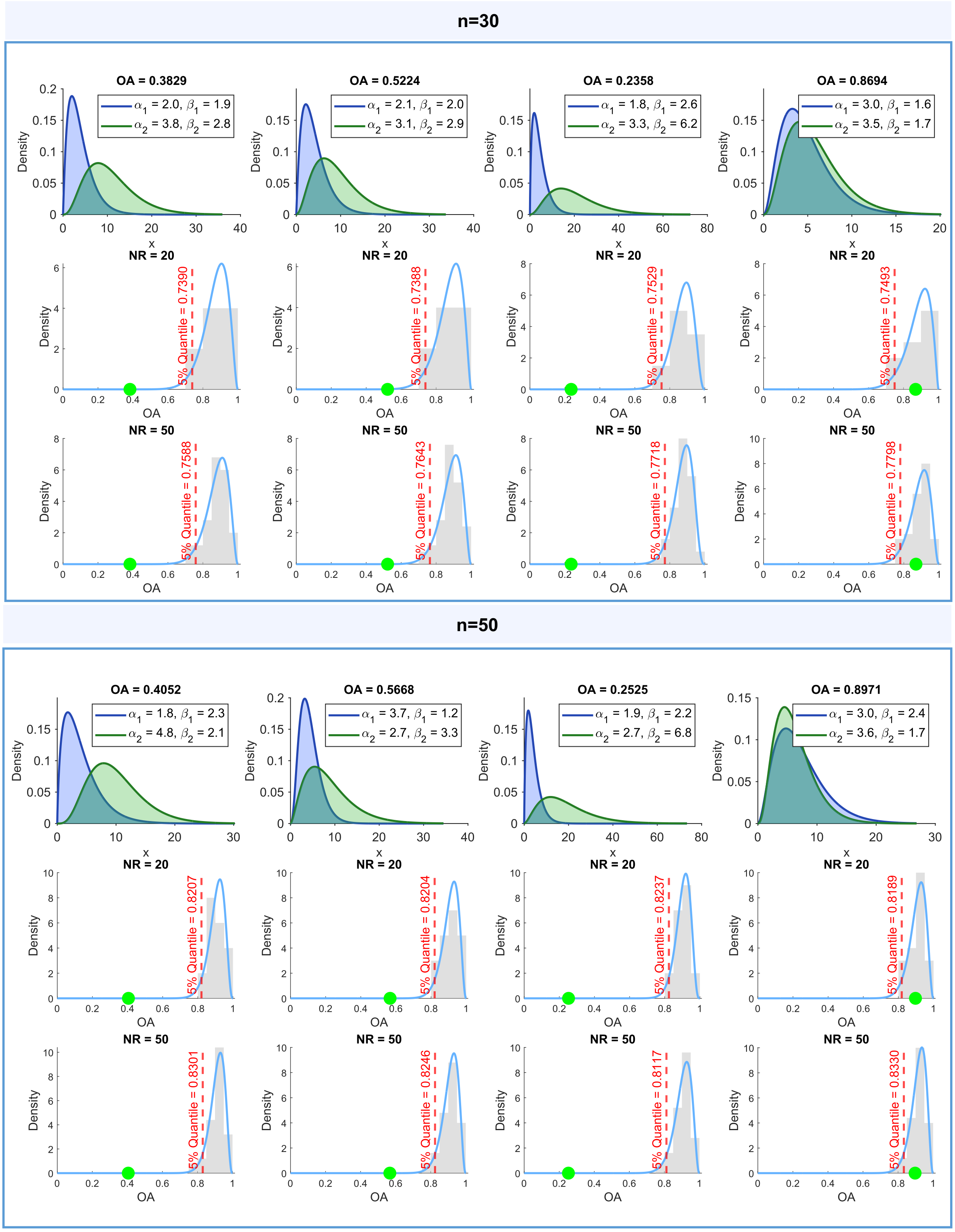


**Figure S1: Simulation results.** The estimated distributions and their overlap for two groups (Group 1, blue curve; Group 2, green curve; the shaded area represents the overlap area) in a single random simulation (top row), along with the sampling distributions of overlap area from $\mathrm{NR} = 20$ (middle row) and $\mathrm{NR} = 50$ (bottom row) based on different cases and a sample size of $n = 10, 20, 30, 50$. Green dots, values of the overlap area; red dashed line, threshold at the 5% significance level.

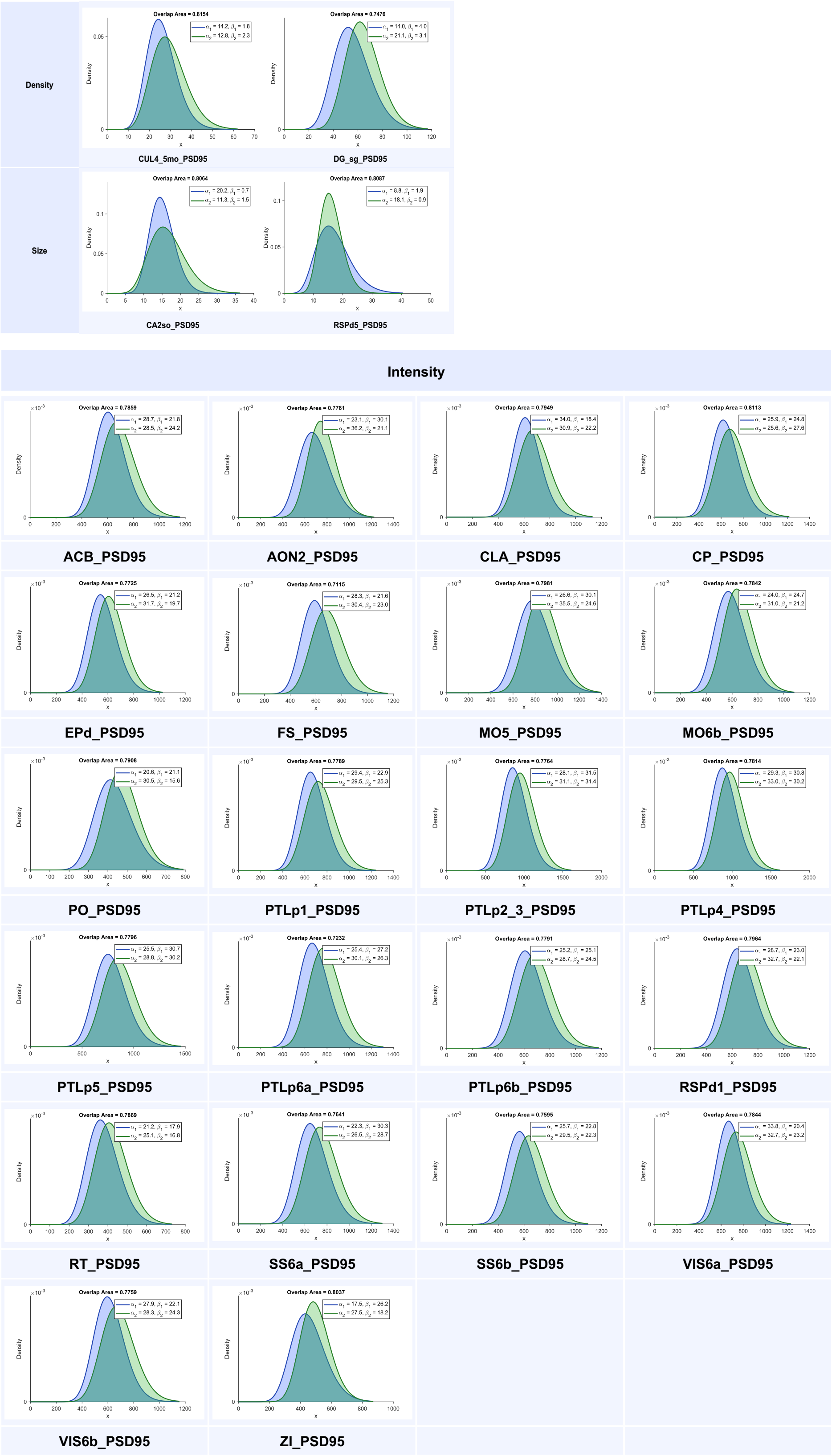


**Figure S2: Real example results.** Population distribution estimates and OA estimates of PSD95 density, size, and intensity in all brain regions showing significant differences.
