## Supplementary material for "The Overlap Area as a Novel Measure of Effect Size in Neuroscience Research": Read_me for App

How to Install and Run overlap_area App in MATLAB
🔧 Installation Steps:
Open MATLAB.

Double-click the overlap_area.mlappinstall file to install the app.

Wait for MATLAB to show the message:
"App installed."

▶️ How to Launch the App:
In MATLAB, go to the top menu bar and click the "Apps" tab.

Scroll down to the "My Apps" section.

Find the icon labeled "app".

Click the icon to launch the app.

🧪 Testing the App:
You can input data for Group 1 and Group 2 to test the overlap area calculation.
